# Highly Distinguished Amino Acid Sequences of 2019-nCoV (Wuhan Coronavirus)

**DOI:** 10.1101/2020.01.31.929497

**Authors:** Jacob Beal, Thomas Mitchell, Daniel Wyschogrod, Jeff Manthey, Adam Clore

## Abstract

Using a method for pathogen screening in DNA synthesis orders, we have identified a number of amino acid sequences that distinguish 2019-nCoV (Wuhan Coronavirus) from all other known viruses in *Coronaviridae*. We find three main regions of unique sequence: two in the 1ab polyprotein QHO60603.1, one in surface glycoprotein QHO60594.1.

## Text

The emerging coronavirus 2019-nCoV(*1*) is of significant world-wide concern as it spreads from its initial point of identification in Wuhan. Identification of significant areas of uniqueness that distinguish such an emerging pathogen may be of value in the development of methods for diagnosis, prevention, or treatment. To this end, we have identified a number of amino acid sequences that distinguish 2019-nCoV from all other known viruses within the family *Coronaviridae*. Amongst these, we find three main regions of unique sequence: two in the 1ab polyprotein QHO60603.1, one in the surface glycoprotein QHO60594.1.

To identify unique sequences, we adapted FAST-NA, a software tool for screening DNA synthesis orders for pathogens(*2,3*) that uses methods for automatic signature generation developed originally for cybersecurity malware detection(*4*). In particular, FAST-NA compares all k-mer sequences of a collection of target sequences to a collection of contrasting sequences in order to identify all k-mer sequences that are unique to the target population. These unique sequences are diagnostic of membership in the population, whereas shared sequences indicate structure that is conserved to some degree.

Here, we applied FAST-NA to identify all of the unique 10-mer sequences in all of the amino acid sequences for 2019-nCoV then available from NCBI: 63 amino acid sequences available in NCBI, comprising a total of 49379 amino acids (*5-8*). For contrasting sequences, we used a July, 2019 snapshot of all protein sequences in family *Coronaviridae* available from NCBI, a total of 50574 sequences comprising a total of approximately 40 million residues. The resulting collection of unique 10-mer amino acids sequences were then concatenated where overlapping within the same parent sequence and trimmed to remove non-unique flanking portions.

All told, this process identifies 61 multi-amino-acid regions as significant unique sequences for 2019-nCoV, comprising a total of 1669 amino acids (3.4% unique and non-repeated), spread across 8 non-duplicative sequences (Appendix Table 1). In addition, we also identified 45 single amino-acid polymorphisms (Appendix Table 2). Figure 1 summarizes the distribution of unique sequence regions across these 8 open reading frame (ORF) sequences. Two of these have notably high amounts of unique content: the large 1ab polyprotein QHO60603.1 has much unique material, though the fraction is not large, while the surface glycoprotein QHO60594.1 has both a large amount and large fraction of unique material.

**Figure 1.**
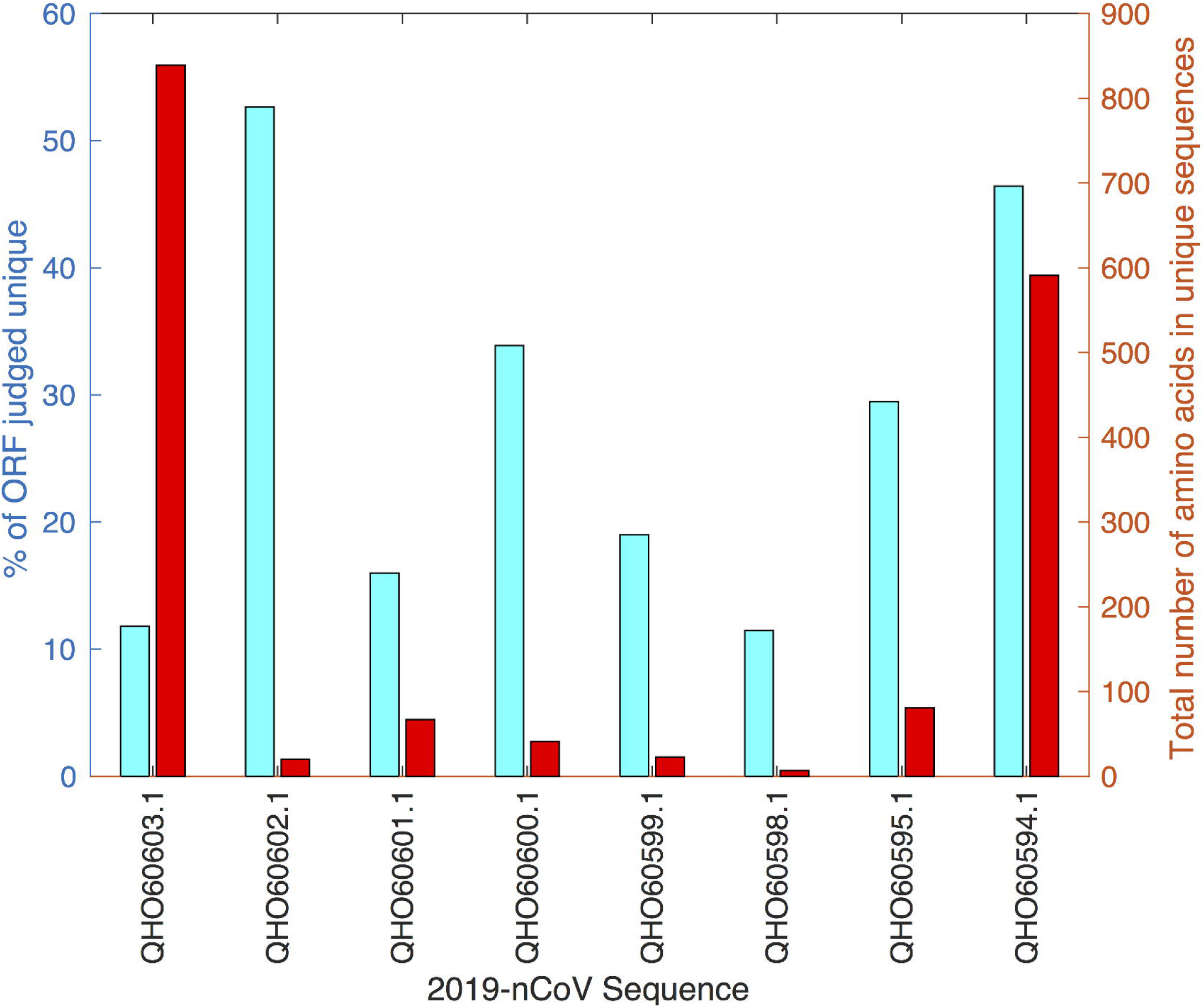
Summary statistics of distinguishing amino acid sequences identified for 2019-nCoV (Wuhan coronavirus), showing the fraction of each ORF judged to be part of unique sequences and the total number of amino acids in unique sequences in the ORF. The large 1ab polyprotein QHO60603.1 has much unique material, though the fraction is not large, while the surface glycoprotein QHO60594.1 has both a large amount and large fraction of unique material.

Further examination shows that the unique material in these two ORFs is strongly clustered. Taking a cluster as any sequence of at least three unique regions with no more than 50 amino acids separating them, we find that QHO60603.1 has two clusters, one spanning from residues 916 – 1294, the other from 6417 – 6715, containing 47% of the unique material in the sequence. The QHO60594.1 sequence, meanwhile, has a single large cluster, spanning from residues 9 to 883 and comprising all of the unique material in the sequence.

In summary, analysis of the amino acid sequences of 2019-nCoV identifies three large highly unique regions of the genome that distinguish it from all other *Coronaviridae*, plus several dozen other smaller regions of uniqueness. We thus hypothesize that these three large regions are likely to be of significance in understanding the evolution and infectivity of 2019-nCoV, in development of countermeasures to mitigate its effects, and in the selection of diagnostic assays to understand and track the origin and spread of this disease, and therefore recommend them as a potential focus of attention.

## Acknowledgments

This research was sponsored by IARPA contract 2018-17110300002 and by the Army Research Office and under Grant Number W911NF-17-2-0092. The views and conclusions contained in this document are those of the authors and should not be interpreted as representing the official policies, either expressed or implied, of the Army Research Office or the U.S. Government. The U.S. Government is authorized to reproduce and distribute reprints for Governmental purposes notwithstanding any copyright notation thereon. This document does not contain technology or technical data controlled under either U.S. International Traffic in Arms Regulation or U.S. Export Administration Regulations.

## Author Bio

Dr. Jacob Beal is a Senior Scientist at Raytheon BBN Technologies, where he leads research on synthetic biology and distributed systems engineering. His work in synthetic biology includes development of methods for calibrated flow cytometry, precision analysis and design of genetic regulatory networks, engineering of biological information processing devices, standards for representation and communication of biological designs, and signature-based detection of pathogenic sequences.

---

**Appendix Table 1.**
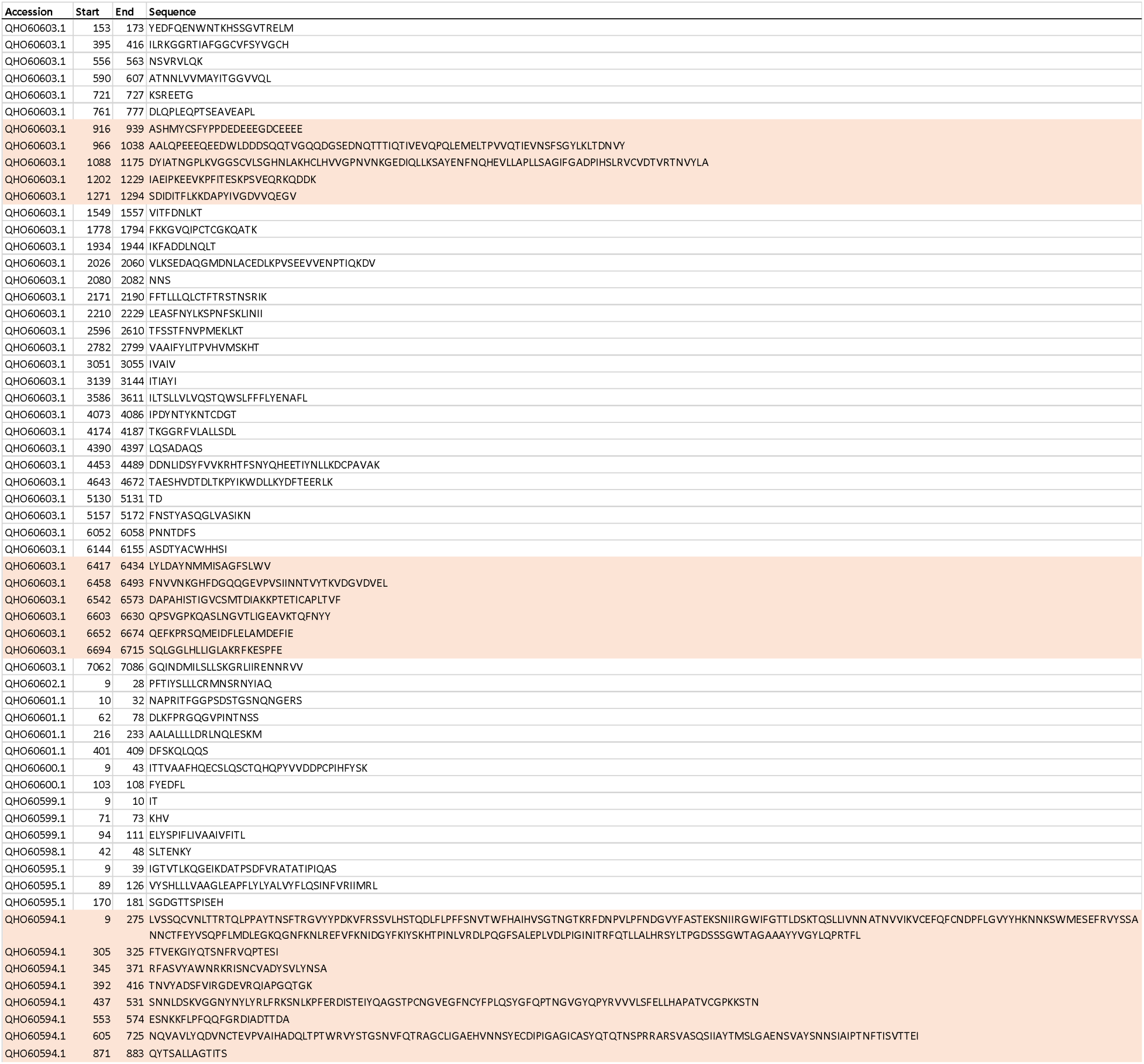
Unique amino acid sequences of 2019-nCoV. Three clusters of unique sequences with less than 50 aa separation are highlighted in red.

**Appendix Table 2.** Additional single-amino-acid polymorphisms of 2019-nCoV.

| Accession | Single AA Polymorphisms location=value |
| --- | --- |
| QHO60603.1 | 37=V 92=E 113=I 279=I 337=K 375=S 444=G 497=A 858=A 1392=V 1439=D 1732=S 1821=T<br>1861=P 1897=N 2006=T 2129=V 2264=G 2452=V 2876=T 3085=L 3668=M 3846=V 3956=F<br>4114=S 4275=A 5038=S 5938=V 6023=E 6100=N 6217=T 6243=A 6298=S 6361=V 6520=V |
| QHO60601.1 | 102=D 127=D 333=T 378=T |
| QHO60600.1 | 64=A |
| QHO60594.1 | 844=A 1083=D 1132=V |
| QHN73809.1 | 3098=L |
| QHD43422.1 | 83=L |

